# Theta oscillations gate the transmission of reliable sequences in the medial entorhinal cortex

**DOI:** 10.1101/545822

**Authors:** Arun Neru, Collins Assisi

## Abstract

Reliable sequential activity of neurons in the entorhinal cortex is necessary to encode spatially guided behavior and memory. In a realistic computational model of a medial entorhinal cortex (MEC) microcircuit, with stellate cells coupled via a network of inhibitory interneurons, we show how intrinsic and network mechanisms interact with theta oscillations to generate reliable outputs. Sensory inputs activate interneurons near their most excitable phase during each theta cycle. As the inputs change, different groups of interneurons are recruited and postsynaptic stellate cells are released from inhibition causing a sequence of rebound spikes. Since the rebound time scale of stellate cells matches theta oscillations, its spikes get relegated to the least excitable phase of theta ensuring that the network encodes only the external drive and ignores recurrent excitation by rebound spikes. In the absence of theta, rebound spikes compete with external inputs and disrupt the sequence that follows. Our simulations concur with experimental data that show, subduing theta oscillations disrupts the spatial periodicity of grid cell receptive fields. Further, the same mechanism where theta modulates the gain of incoming inputs may be used to select between competing sources of input and create transient functionally connected networks.

## Introduction

The entorhinal cortex (EC) acts as a conduit between hippocampal and cortical circuits ^1^. Superficial layers of the EC receive multiple sensory inputs via the perirhinal and the postrhinal cortices and project to all hippocampal subfields ^2^. Given the diversity of inputs that arrive at the EC and its role as a hub, entorhinal networks must necessarily possess two attributes - It must represent an external input reliably and it must have mechanisms that allow it to flexibly select between competing inputs.

Neurons in the medial entorhinal cortex (MEC) translate sensory input into temporally reliable and spatially confined representations ^3–5^. For example, grid cells in layer II of the MEC fire at locations in space that form a striking hexagonally symmetric pattern ^3^. The stability and precision of this pattern is remarkable given many experimentally measured variables in the MEC – inputs to stellate cells and local field potential oscillations – vary noisily as the animal navigates its environment. How can a stable spatial representation be built upon such shaky ground? We discover that the answer lies in the interplay between theta oscillations, a characteristic slow rhythm present in many brain regions, and the intrinsic and network properties of the MEC. Disrupting theta by inactivating the medial septum, a prominent theta generator ^6^, perturbs the spatially periodic receptive fields of grid cells ^7^ and impairs the animal’s ability to navigate ^8^. Each traversal over a particular region in space fails to generate a reliable response. The cumulative effect of this loss of reliability is that the crystalline grid-like structure of the neuron’s spatial receptive field dissipates into an amorphous pattern. In addition to generating a reliable representations, theta also plays a role in transiently coupling different brain regions to form functional networks ^9, 10^. This is evident in lateral entorhinal cortex (LEC) that, in contrast to the allocentric spatial response of the MEC, represents the location of objects in a particular context ^11^, or encode the passage of time between events ^12^. The LEC forms part of a network including the medial prefrontal cortex and the hippocampus that are important in associative learning. Information transfer across the nodes of this network is gated by theta oscillations and the degree of association between different regions is reflected in the degree of theta synchrony between them ^10^.

In a biophysically realistic computational model of an MEC microcircuit we show that a stable grid pattern can be generated by coupling the network to theta oscillations from the medial septum ^6, 13^. Theta oscillations create periodic windows where the network is alternately receptive or resistant to perturbations. If cortical input arrives within the receptive window, it activates the corresponding interneurons. Competitive interactions between interneurons ensure that only those neurons receiving input are active while the others are inhibited. As the input changes, the locus of activity shifts. The rebound properties of stellate cells lead to spikes that mark the transition in activity from one interneuron to another. However, feedback excitation due to these spikes could trigger activity in postsynaptic interneurons that could compete with external inputs. The rebound time scale of stellate cells is such that these spikes are relegated to the least excitable phase of the interneurons. Theta oscillations cyclically order extrinsic inputs, inhibitory spikes by interneurons and rebound spikes by stellate cells. This ensures that the network listens only to extrinsic inputs and ignores its own activity. Further, we argue that the same mechanism - channeling relevant inputs only during the receptive phase of theta while relegating distractors to the resistant phases - is used by multiple brain regions to form transient functionally connected networks.

## Results

### Oscillatory and bistable dynamics of a small MEC network motif

Layer II of the MEC consists of two distinct microcircuits with characteristic patterns of connectivity within and sparse connections across circuits ^1, 14^. Stellate cells and fast-spiking parvalbumin positive interneurons ^15, 16^ form one circuit while pyramidal cells and 5HT3A interneurons form the other ^1, 14^. This disynaptic circuit motif, where principal neurons interact via an inhibitory intermediary, is prevalent throughout the EC ^14, 15, 17^. To understand how a network’s architecture affects its dynamics we simulated a simple network motif (Figure 1b), a building block of the MEC, that consisted of biophysically detailed models of stellate cells ^18, 19^ and inhibitory interneurons ^20^. Stellate cells show subthreshold membrane potential oscillations and generate rebound spikes when released from inhibition ^21^ (Figure 1a). We modeled these properties using a hyperpolarization activated depolarizing current (Ih) ^22, 23^ and an amplifying persistent sodium current (*I*_*NaP*_) ^24^in addition to leak and spiking currents (*I*_*L*_, *I*_*Na*_ and *I*_*K*_). We modeled interneurons using modified sodium and potassium currents that allowed it to spike at high frequency ^20^. In our simulations, both interneurons received supra-threshold input. However, since they inhibit each other, only one of the neurons spiked while the other remained silent. Successive inhibitory spikes from an interneuron activated depolarizing *I*_*h*_ currents in the postsynaptic stellate cell. This eventually drove the stellate cell to spike. Excitatory drive from the stellate cell activated the other interneuron of the pair that, in turn, silenced the first one. The activity of this motif switched rhythmically. The interneurons alternated between episodes of spiking and quiescence, and oscillated out of step with each other. Rebound spikes by stellate cells marked every transition from spiking to quiescence (Figure 1b). When we removed excitation from stellate cells, only one interneuron remained active (Figure 1c). Stellate cells spikes were required to switch the activity of inhibitory interneurons. These transitions can drive other stellate cells to fire, potentially leading to a pattern of activity where different neurons are sequentially activated. As the input to inhibitory interneurons increased so did its spiking frequency ^20^. At higher frequencies, the time between inhibitory spikes was too short for the postsynaptic neuron to fire a rebound spike and cause a switch in the activity pattern (Figure 1d). In this parameter regime the network acted as a bistable switch where one of the interneurons remained active until a transient external perturbation toggled the switch (Figure 1d). When inhibitory input to one of the stellate cells ceased, it emitted a rebound spike that marked the transition from one state of the network to the other (Figure 1d). Thus, depending on the parameter regime, the motif simulated here can act as an autonomous oscillator (Figure 1b) or as a switch (Figure 1d) whose state can be toggled between activity and quiescence by transient external perturbations.

**Figure 1.**
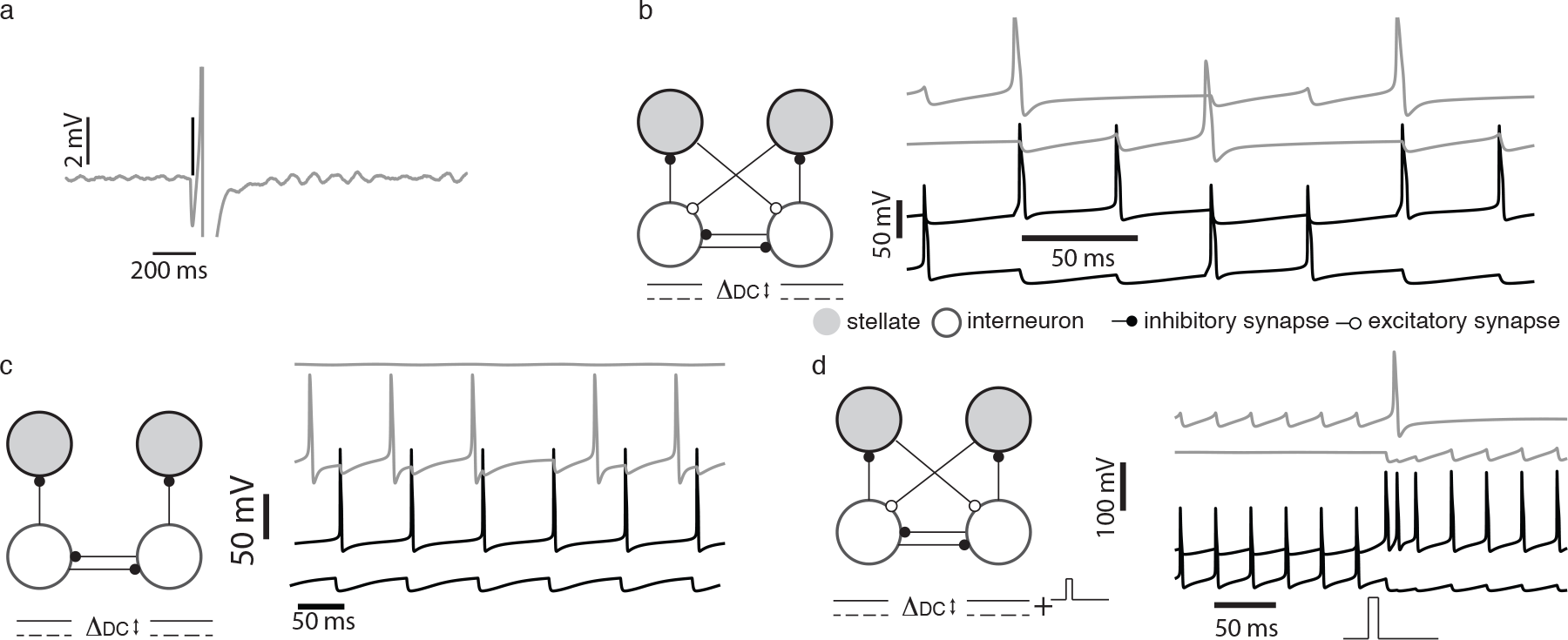
Dynamics of an MEC motif. Gray trace in (a) shows the rebound spiking response of a stellate cell to a single inhibitory spike. The time of the inhibitory spike is shown by the vertical line. In the absence of inhibition, the membrane potential of the stellate cell showed subthreshold oscillations. (b) A supra-threshold input applied to the inhibitory interneurons caused a rhythmic switching of activity in the network motif. Stellate cells spiked (gray traces) when the inhibitory interneuron (black traces) transitioned from activity to silence. (c) Switching failed in the absence of excitatory input; continuous firing of one interneuron (black trace) passively recruited the stellate cell (in gray trace) while the other interneuron (black trace) and corresponding stellate cell remained silent. (d) In a winner-take-all regime both the interneurons received depolarizing inputs higher than that in (b). Only one interneuron fireed continuously, a transient pulse toggled the activity of the inhibitory interneurons (black trace) and stellate cell spiked at the transition (top gray trace).

### Input driven sequences in the MEC network

The EC receives sensory inputs from multiple cortical sources ^1^. Here, we examine the output of a model MEC network in response to a transient input that sequentially stimulates the network. The model consisted of 40 stellate cells and 40 inhibitory interneurons. Input was modeled as a brief pulse that lasted 40ms. After another 85ms, a different, randomly chosen interneuron, was given the same input (Figure 2a). Successive pulses occurred 125ms apart, the period of an 8Hz theta oscillation. All the inhibitory interneurons were connected to each other. Each stellate cell received input from 5 randomly chosen interneurons.

**Figure 2.**
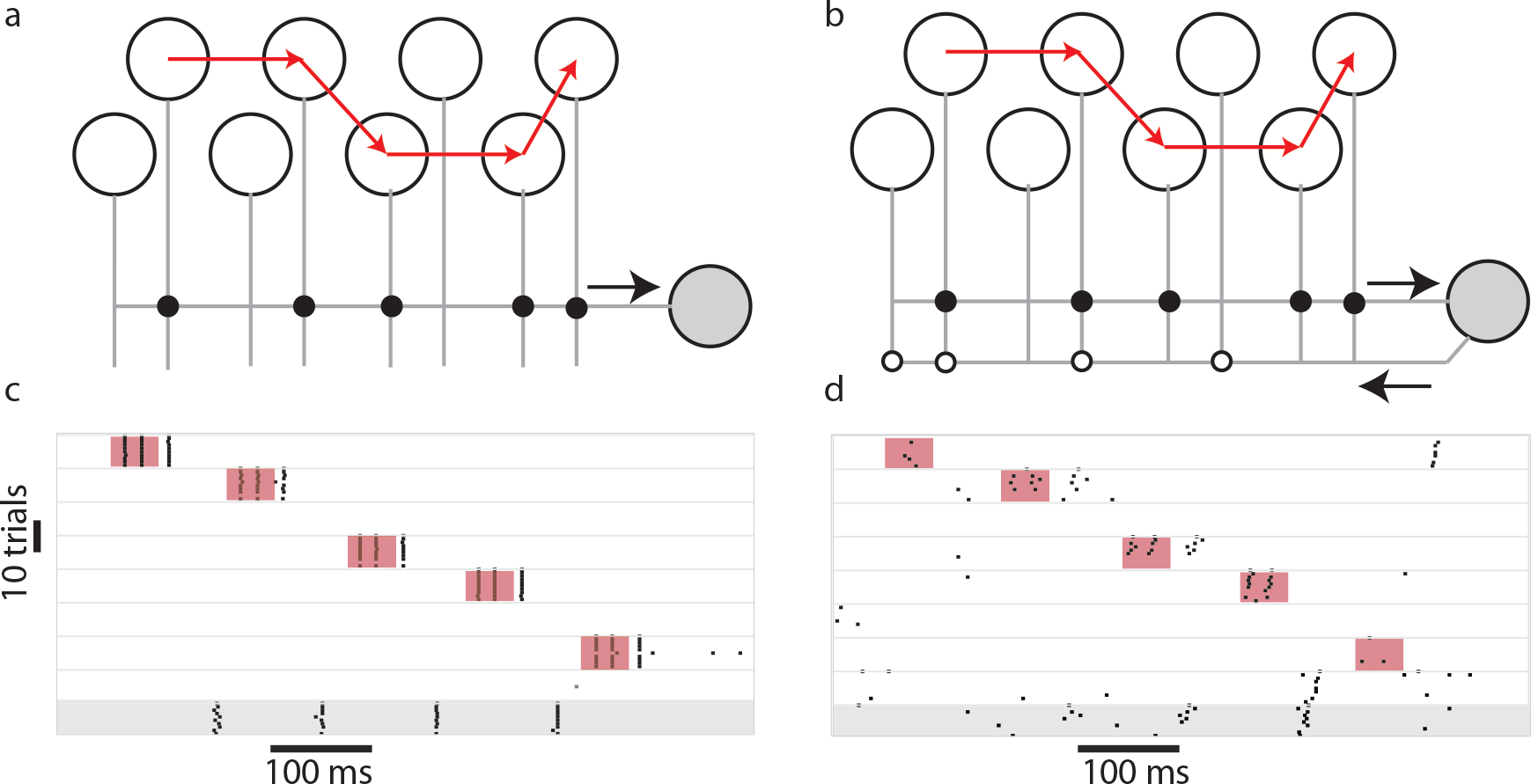
Stability of input driven sequences. (a) Network with a sequential suprathreshold depolarizing pulse (temporal order of the input is shown by the red arrows) driving a subset of interneurons that were connected to a single postsynaptic stellate cell (filled gray circle). The response of the interneurons over 10 trials is shown as a raster plot in (c). The stellate cell (bottom raster shaded in gray) responded reliably over multiple trials. When feedback inhibition from the stellate cell to a randomly selected subset of interneurons (b) was introduced, the response of the interneurons to the sequential input was perturbed and stellate cells did not spike reliably. The duration of the inputs is marked by the colored boxes in (c) and (d)

Our model compared two scenarios, one where there was no recurrent excitation from the stellate cells to the inhibitory interneurons (Figure 2a) and another where each stellate cell randomly connected to 6 interneurons (Figure 2b). In the absence of recurrent excitation from stellate cells, the interneuron that received external input generated spikes that inhibited all the other interneurons. When the pulse moved to a different interneuron, the locus of activity also shifted (Figure 2c) and a rebound spike in postsynaptic stellate cells marked this shift. Since the stellate cell received inputs from multiple inhibitory interneurons, successive shifts from one interneuron to another elicited multiple rebound spikes from the stellate cell (Figure 2c, bottom raster in gray background). Sequential activation of inhibitory interneurons and activation of stellate cells occurred reliably over multiple trials (Figure 2c). Different trials were distinguished by continuous noise trains to stellate cells and interneurons with a mean amplitude that was 10 % of the amplitude of the input to interneurons. When we introduced random excitatory connections from stellate cells to inhibitory interneurons (Figure 2b), the response of the MEC network did not consistently follow the external drive (Figure 2d). Interneurons received competing depolarizing inputs, one from the transient external drive and the other due to stellate cell spikes. The background noise and the history of activation determined which interneuron won this competition and silenced all others. This led to unreliable switching and considerable trial-trial variability in the activity of inhibitory interneurons. Therefore, the firing of stellate cells, that mark the shifts in interneuron activity, was also unreliable across trials (Figure 2d, bottom trace in gray background).

### Theta oscillations reduce trial-trial variability of MEC responses

In a model MEC network, we show that theta oscillations play an important role in generating spatially periodic and temporally reliable sequences of activity akin to that generated by grid cells. Our network consisted of stellate cells and fast spiking inhibitory interneurons. Stellate cells are the most populous cell type comprising nearly 70 % of principal neurons in layer II of the MEC. Of these, one in four are grid cells ^3,25,26^. The receptive fields of different grid cells form an overlapping patchwork that covers space ^3^(Figure 3a). Within a localized region of the MEC, grid cell receptive fields share the same spatial frequency and orientation, but are phase-shifted with respect to each other. We assumed, since stellate cells in our model were activated by inhibition, neurons with overlapping spatial receptive fields that can fire in close temporal proximity, and with similar response patterns, must also receive overlapping inhibitory input (Figure 3b). What are the consequences of this assumption? If receptive field overlap is indeed a reflection of overlap in input connections to stellate cells, it must also constrain the responses of the system in different, independent environments. Environmental and experiential changes to the receptive field of one grid cell must be correlated to similar changes in other cells such that the spatial phase relationships are preserved. Changes in distal environmental cues change the orientation of a grid cell’s receptive field. Changes in the shape of the enclosure can deform the receptive field ^27^. As an animal becomes familiar with the environment, the scale of grid cell receptive fields shrink ^28^. Despite these transformations, the phase relationship between the receptive fields of two grid cells, remains invariant ^29^, suggesting that the spatial contiguity of receptive fields must be a consequence of the topology of the underlying network. Figure 3a shows the overlapping receptive fields of four stellate cells. These receptive fields form a periodic triangular lattice that can be mapped to a torus ^30^. Any straight-line trajectory along a primary grid axis generates a periodically repeating pattern of activity that can be mapped to a circle ^31^(Figure 3a). In keeping with our assumption that overlapping responses are generated by stellate cells with overlapping inputs, we modeled the stellate-interneuron network as a ring where ‘neighboring’ stellate cells that fire in close spatial and temporal proximity were connected to overlapping groups of interneurons (Figure 3b). The strength of synaptic input from an interneuron to the stellate cell layer followed a Gaussian profile. The interneurons were all–to–all connected in this network (Figure 3b). The neighborhood relationship between interneurons was therefore inherited from their connections onto the stellate cell layer. Stellate cells extended random excitatory inputs onto the inhibitory layer. The trajectory of the animal was modeled by sequentially activating neighboring inhibitory interneurons with a transient input. The raster plots (Figure 3c) show the response of a subset of 20 stellate cells to a temporally varying input. Predictably, the response of the network was not reliable across trials and did not always follow the input. The activity of one of the neurons, averaged across 10 trials, is shown in Figure 3e. Due to the ring like topology of the network, the neuron periodically received a supra-threshold input. The neuron was just as likely to fire when an input was present as when it was absent (dashed line in Figure 3e).

**Figure 3.**
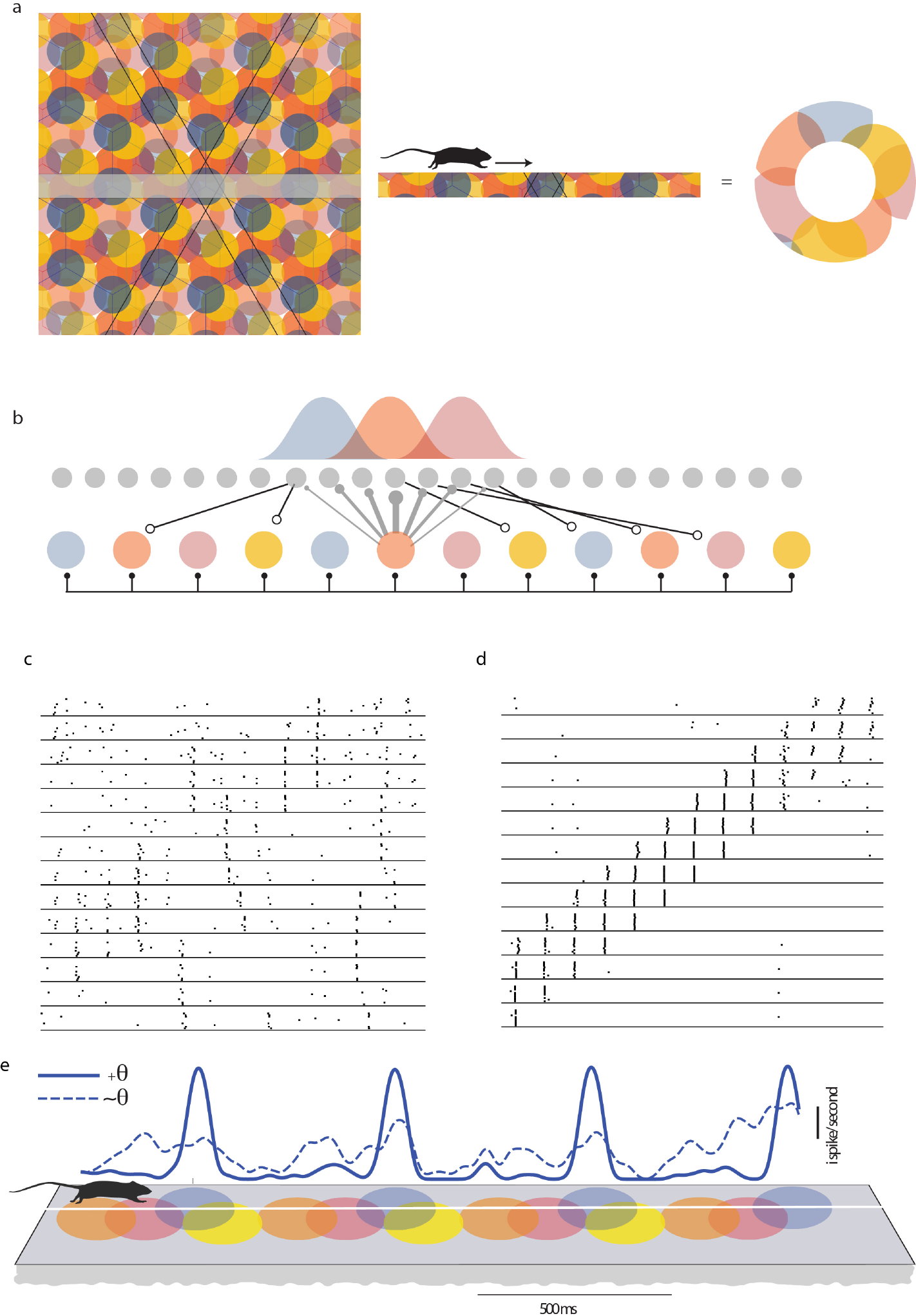
Theta mediated stability of an MEC network. A schematic representation of the overlapping grid-like receptive fields of 4 grid cells is shown in the left panel of (a). As the rat traveled along a primary grid axis (marked in the figure as straight line paths), the grid field repeated periodically (middle panel of (a)). This can be mapped to a circle (right panel of (a)). Stellate cells in the network (gray filled circles in (b)), received inhibitory input from the interneuron layer (coloured circles). The strength of the inhibitory input followed a Gaussian profile (distribution shown in (b)). Each stellate cell connected to a randomly selected group of interneurons. The response of the stellate cells in the absence of theta drive to the inhibitory interneuron layer is shown as a raster plot in (c). When theta oscillations (8Hz) were present, stellate cells evoked a reliable response over ten trials (d). The mean firing rate convolved with a moving Gaussian window is shown in (e) when theta oscillations were present (solid line) or absent (dashed line). The schematic below shows the grid fields of a rat running at a uniform velocity. The temporally periodic spiking pattern of stellate cells appears as a spatially periodic response.

Reliable responses of principal neurons in the EC and the hippocampus are often contingent upon the presence of theta oscillations. One source of theta to the MEC is a central pattern generator in the medial septum ^6^ that extends GABAergic connections to the inhibitory interneurons and periodically modulates its firing rate ^13^. We implemented theta rhythmic modulation of the MEC network by periodically (6-12Hz) driving the entire population of inhibitory interneurons. The pulse like input that was used to drive the neurons remained the same as in earlier simulations (See figure 2 and methods). Neighboring interneurons were recruited in successive cycles of the theta oscillations (125ms apart). We found that the presence of theta oscillations ensured that only those neurons receiving an external input fired and suppressed the activity of all the other interneurons. The output generated in response to a sequential input pattern was stable across noise trials (Figure 3d). The firing rate map (Figure 3e, solid line) calculated for one neuron as an animal traversed a linear track (Figure 3e, schematic) is shown. Assuming that the animal moved at a uniform velocity, it encountered each grid field (alternately, received an external input) after a fixed interval of time and reliably generated a sequence of spikes in response to the input.

### Mechanism of theta induced reliability

How do theta oscillations affect the network such that it responds selectively to an external input and not to the distractors, namely, competing excitatory spikes from stellate cells? The networks simulated in Figure 3 operated in a regime where a transient drive or an excitatory spike can cause a perturbation that toggles the activity of the interneurons (Figure 1c). To understand the effect of theta oscillations on the network, we first simulated a simple network (Figure 4a) where the two interneurons received different constant depolarizing inputs. The neuron receiving the higher input continually spiked while the other neuron was quiescent. In order to toggle the activity of this network, we stimulated the quiescent neuron with a transient pulse. For sufficiently strong pulses, the neuron’s activity switched for the duration of the input (Figure 4b, bottom traces). Weaker pulses, on the other hand, did not evoke a transient switching response (Figure 4b, bottom traces). We then stimulated the interneurons with a periodic theta drive. Theta oscillations modulated the firing rate of the interneurons. Interneurons were maximally depolarized at the trough of theta and were most likely to spike then. When a weak pulse arrived at the quiescent neuron during the depolarizing phase of the theta oscillation, it successfully toggled the network. However, inputs that arrived at other phases did not cause a switch (Figure 4c). Therefore, theta created periodic temporal windows where the activity of the network could be switched from one interneuron to another. In a larger network, we found that when the input arrived at the receptive phases of theta, the stimulated interneurons responded reliably to the input and the locus of activity of the network followed the input (Figure 4d). However, in a large network, merely creating periodic windows where external inputs can drive the network is not sufficient to generate reliable activity. Excitatory spikes from stellate cells can also occur during this receptive phase and perturb the response of the network to an external input. We found that when theta oscillations were present, the stellate cell spikes were tightly synchronized (Figure 4d). These spikes always followed a burst of activity in the presynaptic interneurons (Figure 4d inset, Figure 4f). The interneurons were locked to the trough of theta oscillations while stellate cell spikes occurred at a later phase due to the time taken to generate a rebound spike when released from inhibition. The stellate cell spikes occurred when the interneurons were hyperpolarized and were impervious to feedback excitatory input from stellate cells (Figure 4d,f).

**Figure 4.**
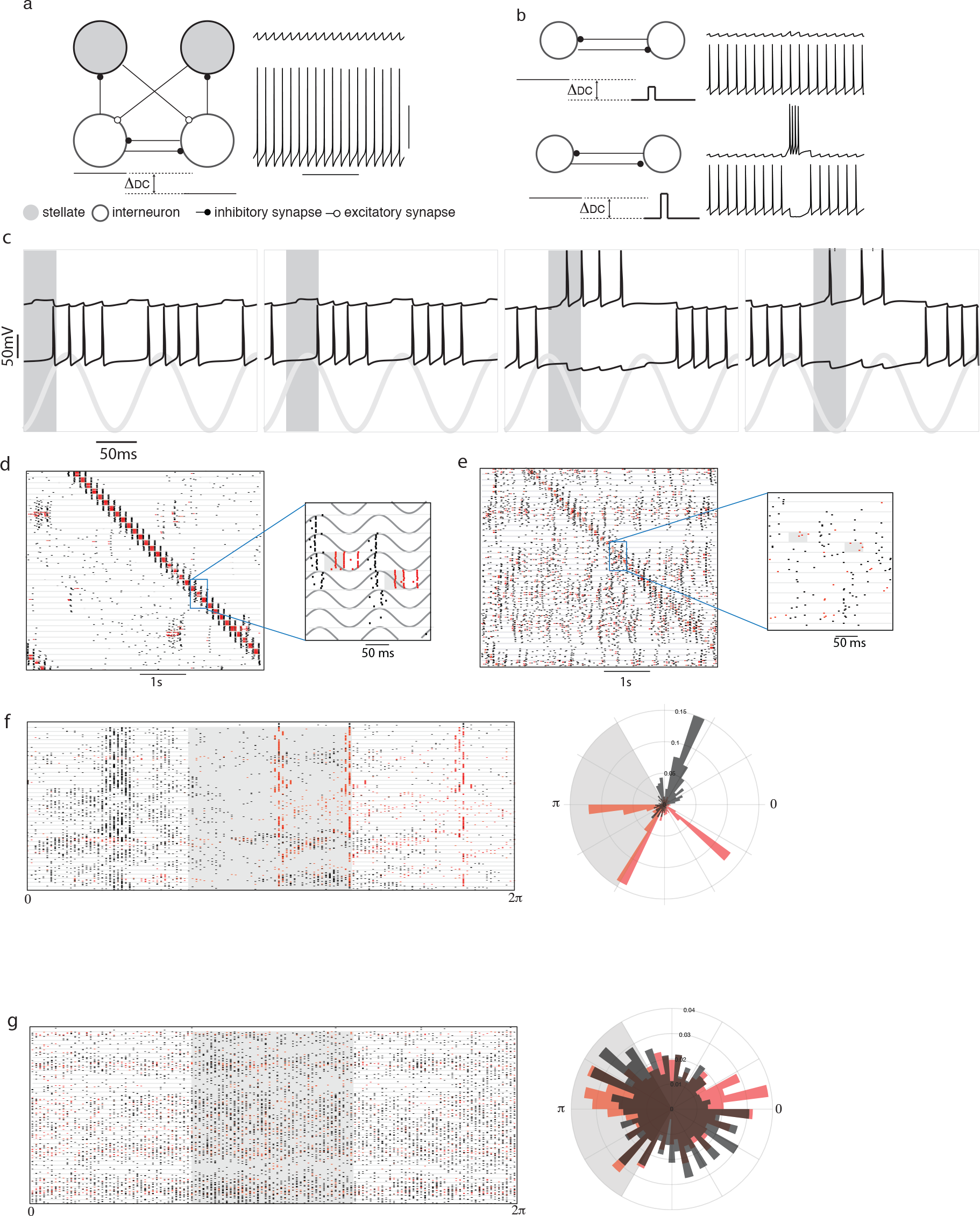
Mechanism of theta induced reliability. (a) Membrane potential of inhibitory interneurons when they received different suprathreshold inputs. The neuron receiving the higher input fired continually (bottom trace) while the other remained silent (top trace). (b) The same network motif as (a) (stellate cells not shown) was simulated. A weak transient pulse was given to the interneuron receiving the lower DC input (top panel of (b)). The stimulated interneuron (top trace) did not switch. A strong pulse caused a successful switch (bottom two traces). (c) Theta rhythmic drive was given to both the interneurons. A weak pulse (duration of the pulse is marked by the gray bar) caused a switch when it occured in some phases of the theta oscillation (bottom gray trace) but not during others. (d) raster plot showing the reliable response of a subset of 20 interneurons (red line) and 20 stellate cells (black lines) from a larger network of 80 neurons, to an external input (marked as a gray bar in the magnified raster plots). The response of the same network in the absence of theta oscillations is shown in (e). Each row in d and e shows the response of a neuron. 10 trials are plotted between the horizontal lines in the raster plots (f) Left. The raster plot shows the phase (with respect to the theta oscillation) at which stellate cells (black lines) and interneurons (red lines) spiked. The gray area marks the phase of the oscillation when the stimulus was present. Right. Polar plot showing a histogram of the phases where spikes occurred. The external input is marked in gray. (g) shows the response of the network when theta oscillations were absent. The phase was defined in terms of the input pulse that sequentially and periodically stimulated neighbouring interneurons.

The input triggered response of the network revealed the segregation of time between the inhibitory interneuron spikes and the stellate cell responses following a transient input (Figure 4d). When theta was present and the external input occurred at a receptive phase, it was followed by a reliable burst of interneuron spikes, followed by the activity of stellate cells. In the absence of theta also, stellate cells fired after each burst of interneuron spikes (Figure 4e and the inset). However, the stellate cell spikes were broadly distributed throughout the time between successive inputs to the network (Figure 4e and Figure 4g). Thus, stellate cells could effectively perturb the activity of the network in a manner that derailed its response to an external drive.

### Theta induced reliability persists over a range of frequencies

Theta oscillations span a range of frequencies from 6 to 12 Hz in rodents. The frequency increases linearly with an increase in the animal’s velocity ^32^. As the animal’s velocity increases, it encounters grid fields more rapidly. This translated in our model as an increased rate at which external pulses arrived at neighboring interneurons. We found that the network responded reliably to external inputs for a range of theta frequencies (Figure 5c). We calculated the reliability across trials measured as the pairwise dissimilarity between spike trains (see methods). This measure showed greater reliability for theta between 7-12 Hz compared to a network that was not driven by theta oscillations (compare between circles and triangle points in the box plot Figure 5c). The upper bound on the theta frequency that led to a reliable response was determined by the time scale of the rebound kinetics of stellate cells. Input that arrived near the trough of theta caused a burst of spikes in inhibitory interneurons that, in turn led to rebound spikes in postsynaptic stellate cells. As the frequency of theta oscillations increased, the time difference between the inhibitory bursts and the rebound spikes remained nearly the same. However, the phase at which stellate cell spikes occurred started to shift towards the trough of the theta oscillation cycle and competed with external input pulses. In the network this appeared as a shift in the mean phase of stellate cell spikes (Figure 5a,b). Further, the standard deviation of the spike time distribution increased such that more spikes occurred during the receptive phase of the theta oscillation (Figure 5a, b) and disrupted the ability of the network to follow an external input.

**Figure 5.**
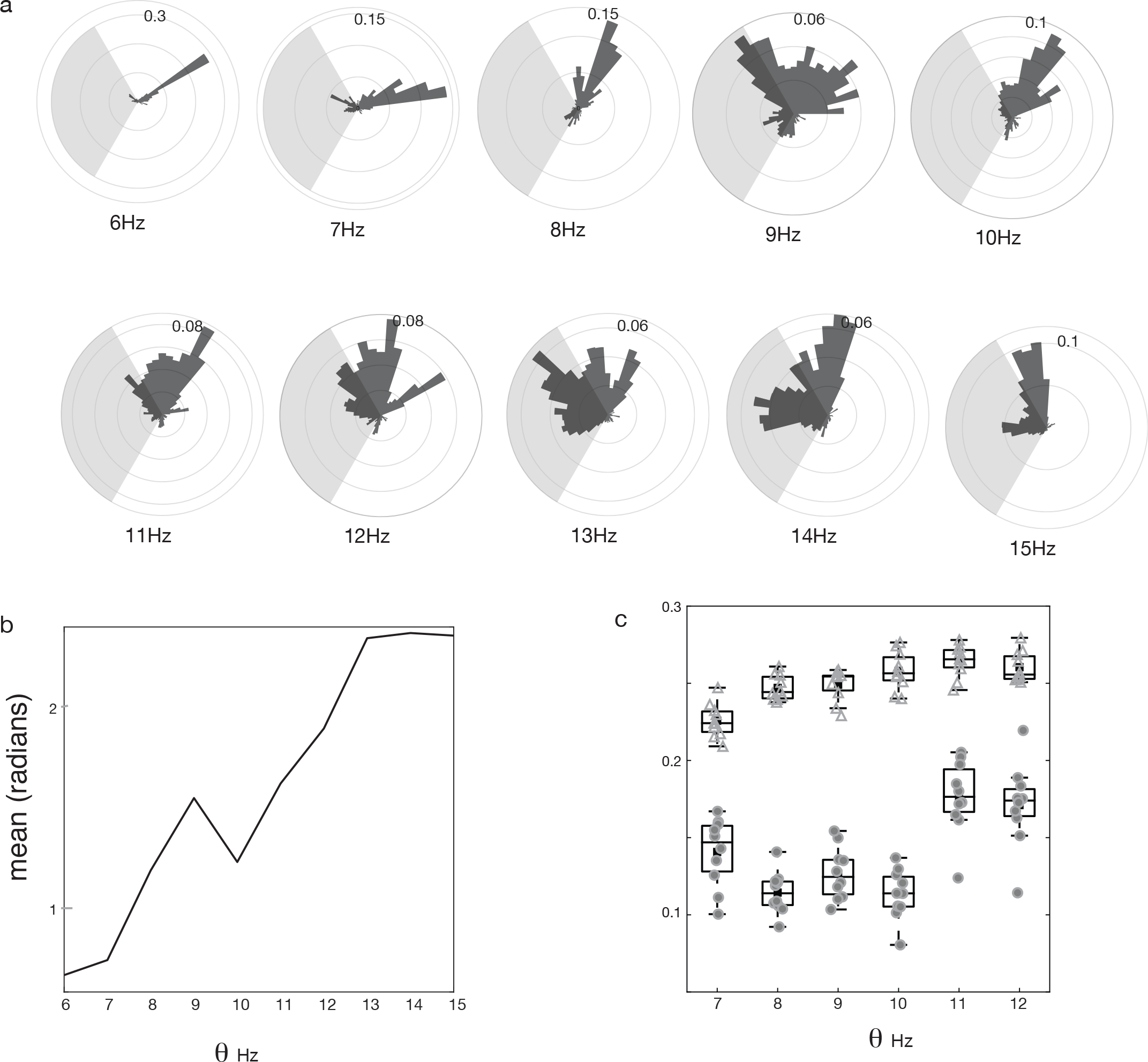
Reliability across a range of theta frequencies. (a) Polar plots showing the distribution of phases of stellate cell spikes for a range of theta frequencies from 6Hz to 15Hz. (b) The circular mean of the phase distribution of spikes plotted against theta frequency (c) box plot of the spike distance in the presence (filled circles) and absence (triangles) of theta as a function of the rate at which input pulses stimulated successive interneurons.

### Theta oscillations gate the transmission of competing inputs

When theta oscillations are present the network reliably follows external inputs that arrive at the receptive phase of theta. Here we examine the response of the network when competing external inputs occur at different phases of the theta cycle. We simulated a network consisting of two groups of interneurons connected to the same pool of stellate cells. All the interneurons were coupled to each other. Each group of interneurons extended inhibitory connections to the stellate cells forming two separate ring networks (Figure 6a). The upper ring of interneurons received a transient pulse that traveled in a counter-clockwise direction while the lower ring received an input that traveled clockwise. The two input streams, clockwise and counterclockwise, competed to elicit a response in the same pool of stellate cells. The amplitude of both the inputs were kept constant while their phase relationship with respect to an external common theta oscillation was varied. In the first of the cases tested (Figure 6b), one of the inputs (clockwise) arrived when the interneurons were near their most hyperpolarized phase while the second input (counterclockwise) arrived at the depolarizing phase. Predictably, the second input succeeded in eliciting a sequence of spikes in the lower ring of interneurons that entrained the stellate cells and inhibited all the other neurons. We then varied the phase of the clockwise input such that the pulse occurred progressively closer to the phase of the counterclockwise input (Figures 6b, Figure 6d and Figure 6e). When the input crossed a particular phase of the theta oscillation (Figure 6b bottom row and 6e), it elicited spikes in inhibitory interneurons of the lower ring. This inhibited all the other interneurons including the neurons on the upper ring of the network. Therefore, even though the counterclockwise inputs occurred during a depolarized phase of theta, the response of the network followed the clockwise inputs because it occurred earlier during the theta cycle. Thus, the mechanism whereby the network selects between inputs is determined, not merely by coherent theta gain modulation of inputs, but also by the temporal order and the phase at which the inputs occur.

**Figure 6.**
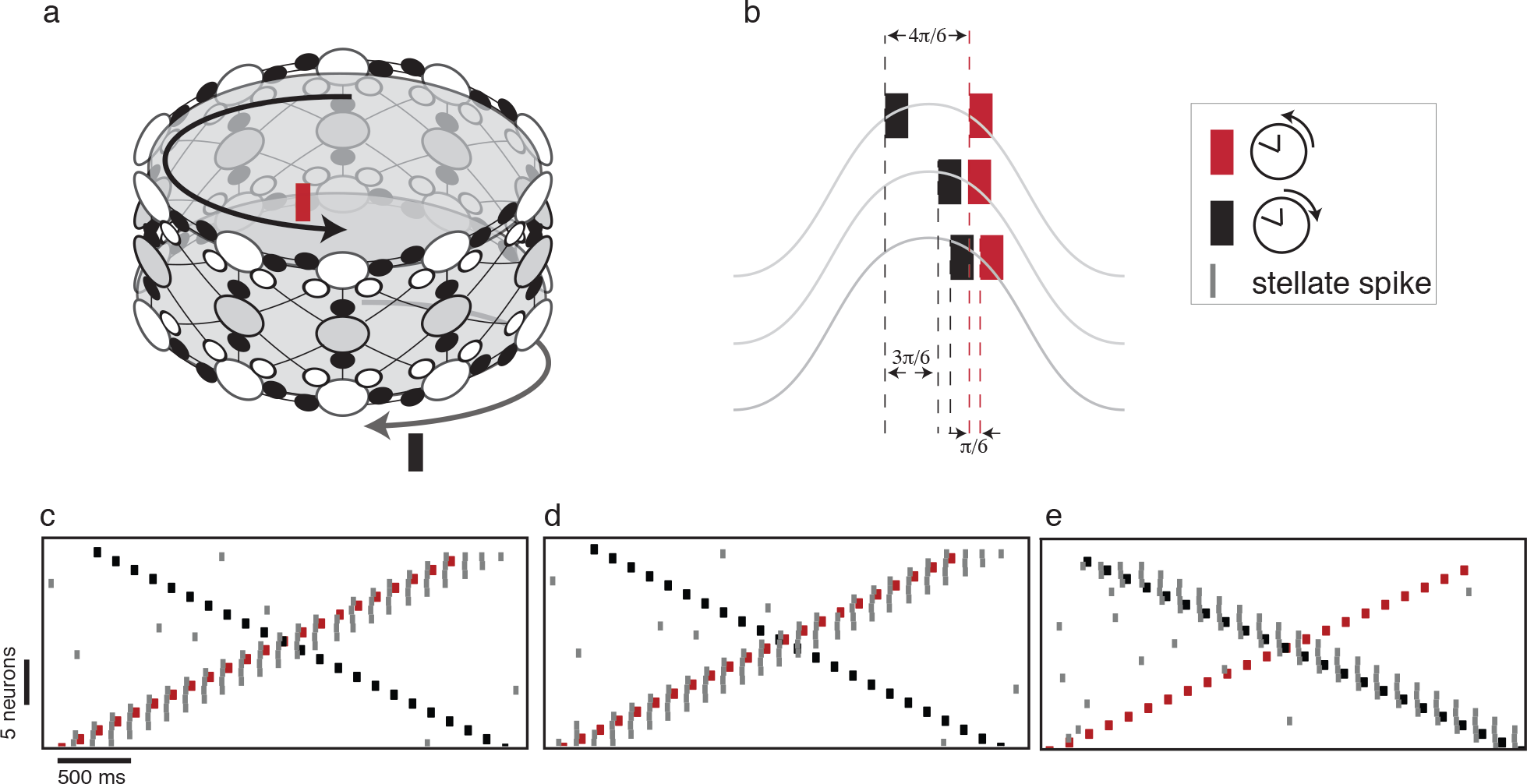
Theta gates transmission of competing inputs. (a) Topology of the network. Two sets of interneurons arranged on different rings (top and bottom empty circles) inhibit the same stellate cell population (middle ring with filled circles). Two seperate input pulse trains are given to the two inhibitory population. The input to upper ring followed a counterclockwise activity pattern while input to the lower ring followed a clockwise sequence. (b) The phase of each input within a single cycle of theta for three different cases. The response of the network to these patterns of input are shown in (c), (d) and (e). The peak of the oscillation corresponds to the maximally hyperpolarized phase of theta (c) The input arriving at the depolarizing phase (red) elicited a spike (gray line) in the post-synaptic stellate cell while the one arriving during the hyperpolarized phase elicited none (this case corresponds to the top trace in (b)) (d) Both the inputs arrive at the depolarizing phase. The input that caused a maximum depolarization (red) elicited a successful response (this case corresponds to the middle trace in (b)). (e) The inputs to both the clockwise and the counter-clockwise rings were shifted. This switched the stellate cell from following the counter-clockwise input to following the clockwise input.

## Discussion

We showed that the MEC interneurons are receptive to external inputs only within cyclic windows defined by theta oscillations. Further, theta oscillations corralled stellate cells to spike synchronously at a phase where inhibitory interneurons were hyperpolarized and less receptive to excitatory inputs. This prevented stellate spikes from depolarizing randomly connected postsynaptic inhibitory interneurons that could compete with external inputs to other neurons. This mechanism to generate reliable sequences can also be harnessed to ensure that the entorhinal cortex selectively gates inputs such that some inputs that arrive at specific phases are transmitted to postsynaptic targets while others are blocked.

Our model suggests that stable sequential activity is contingent upon the presence of theta oscillations. However, recordings from Egyptian fruit bats show that grid like receptive fields can be formed in the absence of continuous theta oscillations ^33^. This seems at odds with our model and experiments in rodent MEC where excising theta reversibly perturbs the grid like structure of the receptive fields of MEC neurons ^7^. One way to reconcile these contradictory observations is to assume that the MEC network in bats receives large amplitude inputs. This can lead to a stable response even in the absence of theta oscillations (Figure 4b, bottom traces). However, given that the MEC is a hub that receives multiple inputs, relying only on the amplitude of the input would impair its ability to selectively respond to some inputs while ignoring others. An additional layer of control can multiplex between similar inputs. Multiplexing is typically implemented using oscillations ^34, 35^. We used theta oscillations to modulate the gain and amplify inputs arriving at some phases while attenuating the effect of others. This phase dependent gain modulation does not require a continuous fixed frequency oscillation. Interestingly, even though bats lack a persistent oscillatory signal like theta in rats, they generate fluctuating low frequency local field potentials to which a significant proportion of principal cells are locked ^36^. Can our model network use these low frequency inputs to generate a reliable output? We showed that reliability of our model network responses progressively deteriorates in the high theta frequency regime (> 11Hz) (Figure 5c). However reliable responses to low frequency oscillations are still possible. At low frequencies, the interneurons fire over a longer duration corresponding to a widened window of depolarization. The hyperpolarizing phase of the theta oscillation tends to shut the response of the interneurons and triggers a rebound excitation in stellate cells (Figure 7a). Stellate cell spikes continue to occur during the hyperpolarizing phase of theta oscillations (Figure 7b) and do not perturb the inhibitory network. Bats show a large variability in the frequency of local field potential fluctuations unlike theta oscillations in rodents that vary over a smaller range. In our model network the input driven switch from one interneuron to another is rapid due to inhibitory competition between fast-spiking interneurons and occurs within a single theta cycle. Therefore, the network activity can respond to fluctuations in a cycle-by-cycle manner, relegating distractors to a hyperpolarized phase in spite of variations in instantaneous frequencies. Changes in instantaneous frequency that occur on a slower time scale than the switching time scale have little impact on perturbing the input driven dynamics of the network.

**Figure 7.**
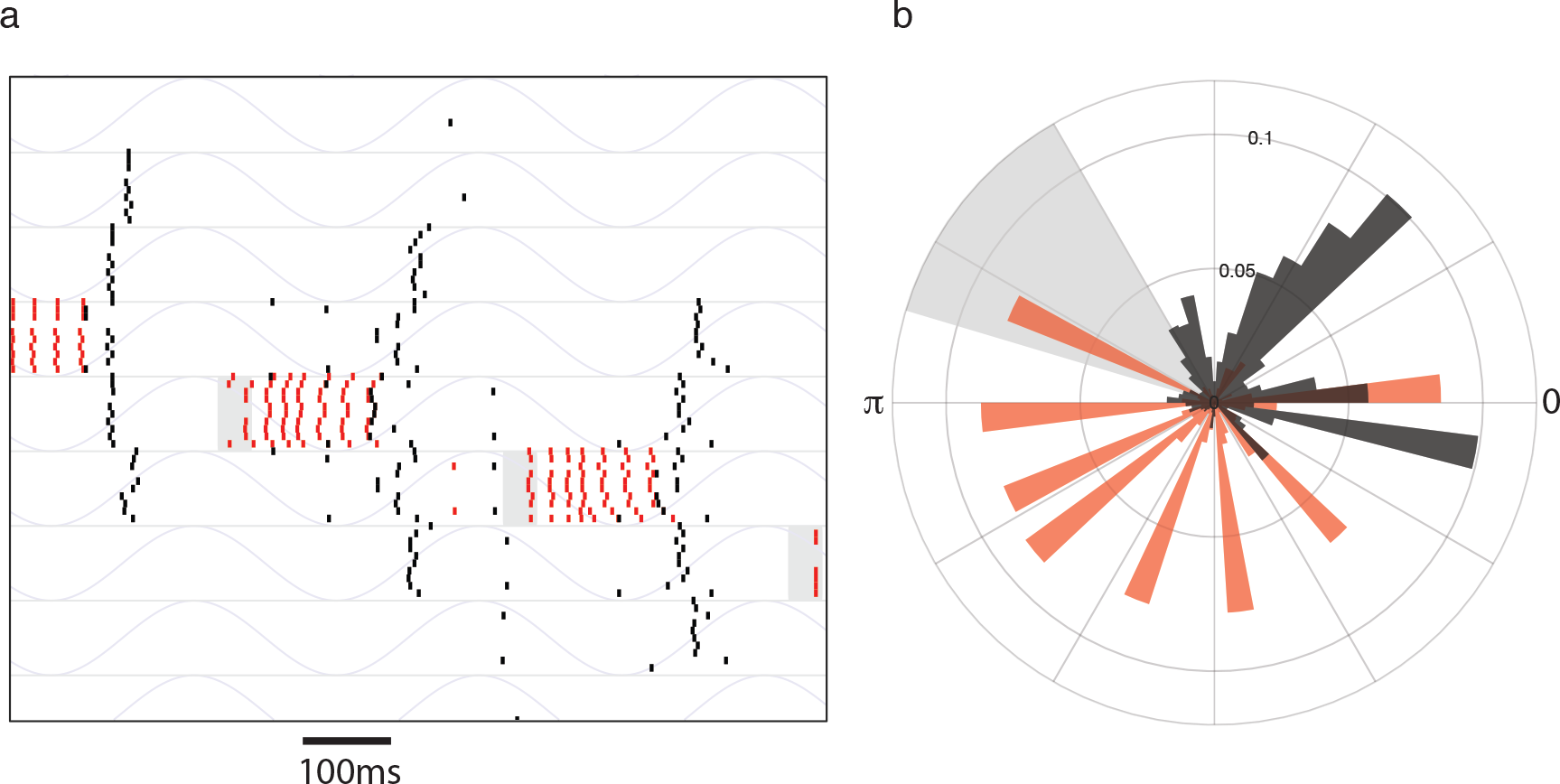
Response of the network to a slow oscillation (2 Hz). The input (shown in gray) toggles the activity of interneurons that continue to generate a burst of spikes (a). Rebound spikes by stellate cells occured at a phase that did not perturb the sequence. (b) histogram of the phase at which the stellate cells (dark bars) and the inhibitory interneurons (red bars) generate spikes.

Our model demonstrated that theta oscillations can gate the transmission of information between different brain regions. One can selectively couple two regions by ensuring that the phase of theta is coherent across these regions and information is transmitted during a restricted phase window of theta. Inter-regional interactions via oscillatory phase coherence is not restricted to circuits including MEC, but widespread across many cortical and subcortical structures ^37–40^. A number of behaviors depend on recruiting a broad network of regions. For example, hippocampal and amygdalar circuits show theta coherence when animals are exposed to anxiety inducing situations ^41^. Working memory in rodents ^42^ recruits hippocampal and medial prefrontal cortex via theta synchrony. Many adaptive behaviours coincide with an enhanced coherence in the phase of the local field oscillations between different brain regions ^?, 43^. Memories at various stages of encoding, consolidation and retrieval invoke different configurations of brain regions that are dynamically assembled by theta phase coherence across these regions. During the early phases of learning an association between a conditioned and an unconditioned stimulus, the hippocampal - lLEC coupling is characterized by phase synchronized theta oscillations. As learning progresses, the phase synchrony between the hippocampus and LEC decreases with a concomitant increase in LEC-medial prefrontal cortex synchrony ^10^. Theta synchrony is a read-out of increased information transfer across brain regions. However, finer control over the input during each theta cycle is required to ensure that MEC networks selectively listen to or ignore incoming inputs. What are the mechanisms that ensure the right inputs arrive at the right phase of theta? Inhibitory interneurons that participate in generating and maintaining hippocampal theta rhythms broadcast rhythmic inhibition that targets inhibitory interneuronsin other areas ^13^. These, in turn, synchronize principal neurons. The effectiveness of inhibition onto principal neurons can alter the degree of synchronization and the phase of principal neuron spikes. In the hippocampal-medial prefrontal cortex circuit, this is likely controlled by neuromodulators like dopamine ^9^ that can effectively shift the phase of principal neuron spikes with respect to a theta oscillation.

Our simulations operated in a regime where the system responded to a sequential external drive that acted as a toggle that moved the locus of activity from one neuron to another. Stellate cells spiked and registered the temporal location of this transition. In the MEC network, GABAergic connections onto the principal neurons show spike timing dependent plasticity that can enhance the weights of inhibitory connections for those synapses where the postsynaptic stellate cell spikes after the inhibitory interneuron ^44^. Repeated sequential activation of the same network will therefore lead to changes in synaptic weight that introduce asymmetries in the network architecture ^45^. These asymmetries can influence the autonomous activity of the network (Figure 1b) to follow sequences that were reinforced by past inputs. These sequences may appear as episodes where the activity of the network is replayed ^46^in the absence of any external inputs.

In sum, our study highlights the central role of theta oscillations in generating reliable sequences and forming transient functionally connected networks. This is possible due to the fortuitous similarity of the timescales of theta oscillations with the intrinsic conductances of neurons and the network architecture of the MEC.

## Methods

The MEC circuit was modeled as a network of stellate cells and inhibitory interneurons. To explain the mechanism of switching and the role of theta oscillations in generating reliable sequences, we used two network motifs: one with two stellate cells and two interneurons (Figures 1 and Figure 4a,b); and the other consisting of 40 stellate cell and 40 interneuron (Figures 3, 4d-g and 5, 7). To explain the role of theta in choosing among the competing external inputs (Figure 6), we simulated a network of 40 stellate cells and 80 interneurons.

### Neuron Models

Stellate cells and interneurons were modeled as conductance based, single compartment spiking neuron models. Stellate cells were endowed with voltage dependent ion channels that were active at subthreshold membrane potentials, apart from sodium, potassium and leak ion channels involved in spike generation. These ion channels enabled the stellate cell to produce sustained oscillations in response to a constant depolarization ^18, 19, 22^.

The current balance equation for the equivalent circuit model of the membrane is given by,

### Stellate cell

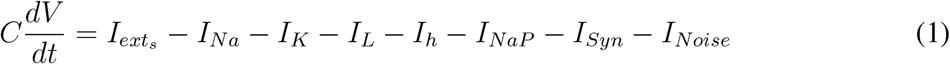

### Interneuron

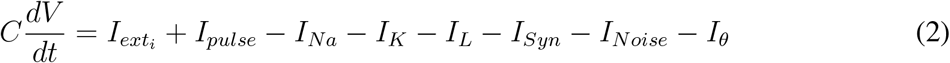

Ionic currents were modeled as follows,

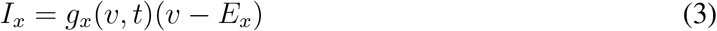

where *g*_*x*_(*v*, *t*) = maximal conductance × *f*(state of gating variables). The gating variables followed first order kinetics. For example, the gating variable, *m* was given by,

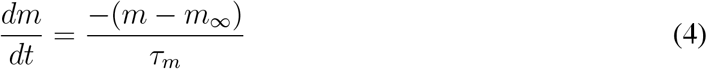

where both *m*_∞_ and *τ*_*m*_ are functions of voltage.

An equivalent representation is

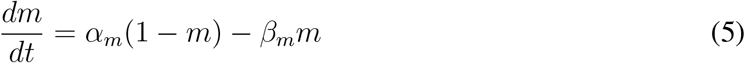

with *m*_∞_ = *α*_*m*_/(*α*_*m*_ + *β*_*m*_); *τ*_*m*_ = 1/(*y*_*α*_ + *τ*_*m*_)

The functional form of each ionic current and the gating variables, maximal conductances and reversal potentials are provided in the tables below.

### Synapse Model

Stellate cells were randomly connected to inhibitory interneurons with an excitatory synapse. Interneurons sent inhibitory connections to stellate cells as well as to other interneurons. The inhibitory population was modeled as an all-to-all connected network. The synapse was modeled as conductance with a gating variable modulated by the presynaptic voltage.

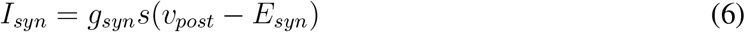

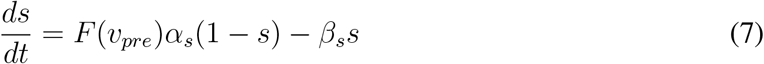

where, *F* (*v*_*pre*_) = (1 + tanh (*v*_*pre*_/4))/2, models the opening of a synaptic ion channel in response to action potential generated by the presynaptic neuron. The reversal potential determined the nature of synapse to be excitatory (when set to 0mV), or inhibitory (when set to −80mV).

### Connectivity

We modeled three network types - A small network motif comprised of two stellate cells and two interneurons (Figure 1b), and two larger networks with stellate cells and interneurons arranged on rings. In one of the larger networks (Figure 3b) with 40 stellate cells and 40 interneurons, the population of stellate cells and the population of interneurons were arranged on separate rings. Each interneuron sent projections to five adjacent stellate cells while each stellate cell sent projections to six randomly chosen inhibitory interneurons. In the network simulated in Figure 6, 80 interneurons were divided into two sub-populations 40 interneurons each. Each sub-population connected to the common pool of stellate cells. The neighboring interneurons on each ring connected to neighboring stellate cells on the stellate cell ring.

### External input to the network

Theta rhythmic input to the nework was provided by periodically modulating the excitability of interneurons using a sinusoidally varying conductance,

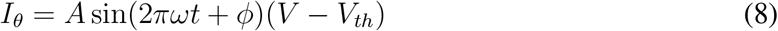

where, *A* is the amplitude of theta rhythm; *ω* its frequency in Hz. *V*_*th*_ is the threshold voltage for the theta drive.

In addition to a constant depolarizing input, a sequential pulse like input was used to drive the inhibitory interneurons. The temporal profile of the transient external pulse for an interneuron *i* is given by,

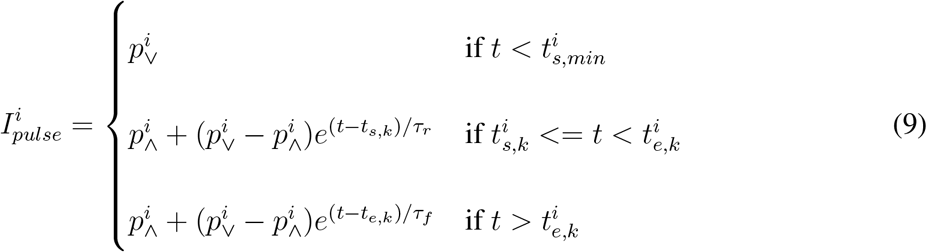

The pulse rises from a baseline value, 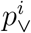, at time, 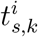, to a maximum, 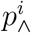, with a rise time of *τ*_*r*_. At 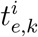 the pulse is switched off and falls to the baseline value with a fall time, *τ*_*f*_. The index *k* indicates the *k*^*th*^ pulse to the neuron. Successive pulse-like inputs were then given to a neighbouring interneurons in successive theta cycles. The start time of a pulse to the *i*^*th*^ neuron was calculated using the following prescription,

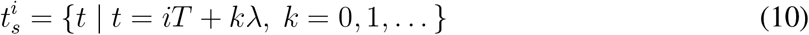

*T*, the inter-pulse time interval matches the period of the theta oscillation. λ is the inter-pulse time interval for a single interneuron. In the ring networks simulated here, the pulse visits all the interneurons before arriving back at the same neuron. Therefore we set λ = *NT*. The duration of the pulse was set to *p*_*width*_. The end time, 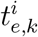, of a *k*^*th*^ pulse to neuron *i* is,

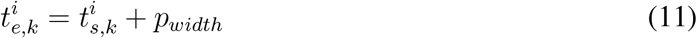

A similar sequence of pulses were given to the two rings of interneurons in Figure 6a. The top ring received inputs in a counter-clockwise direction while the order of inputs to the bottom ring followed a clockwise direction.

### Measure of reliablity

To compare the reliability of the responses of the network for a given sequence of inputs across noisy trials, we used a measure of smilarity between spike trains termed SPIKE-distance ^47^. This measure can be calculated for each neuron across all pairs of trials and averaged over time. This value was calculated for a sub-set of neurons (*N* = 8) that received input for all the theta frequencies that were simulated. The distribution of mean reliability across trial for each neuron is shown in Figure 5c. The analysis were implemented using a Python library, PySpike ^48^.

All the simulations were performed using home-grown C++ library, insilico, that uses odeint, a boost C++ library to solve ordinary differential equations. The differential equations were integrated using an euler method with the time step of 0.01ms. The codes for running the simulations and analysing the outputs are available in the github repository https://github.com/arunneru/theta gates reliable sequences mEC.

The odeint library can be found at the following website: http://headmyshoulder.github.io/odeint-v2/.

**Table 1:**
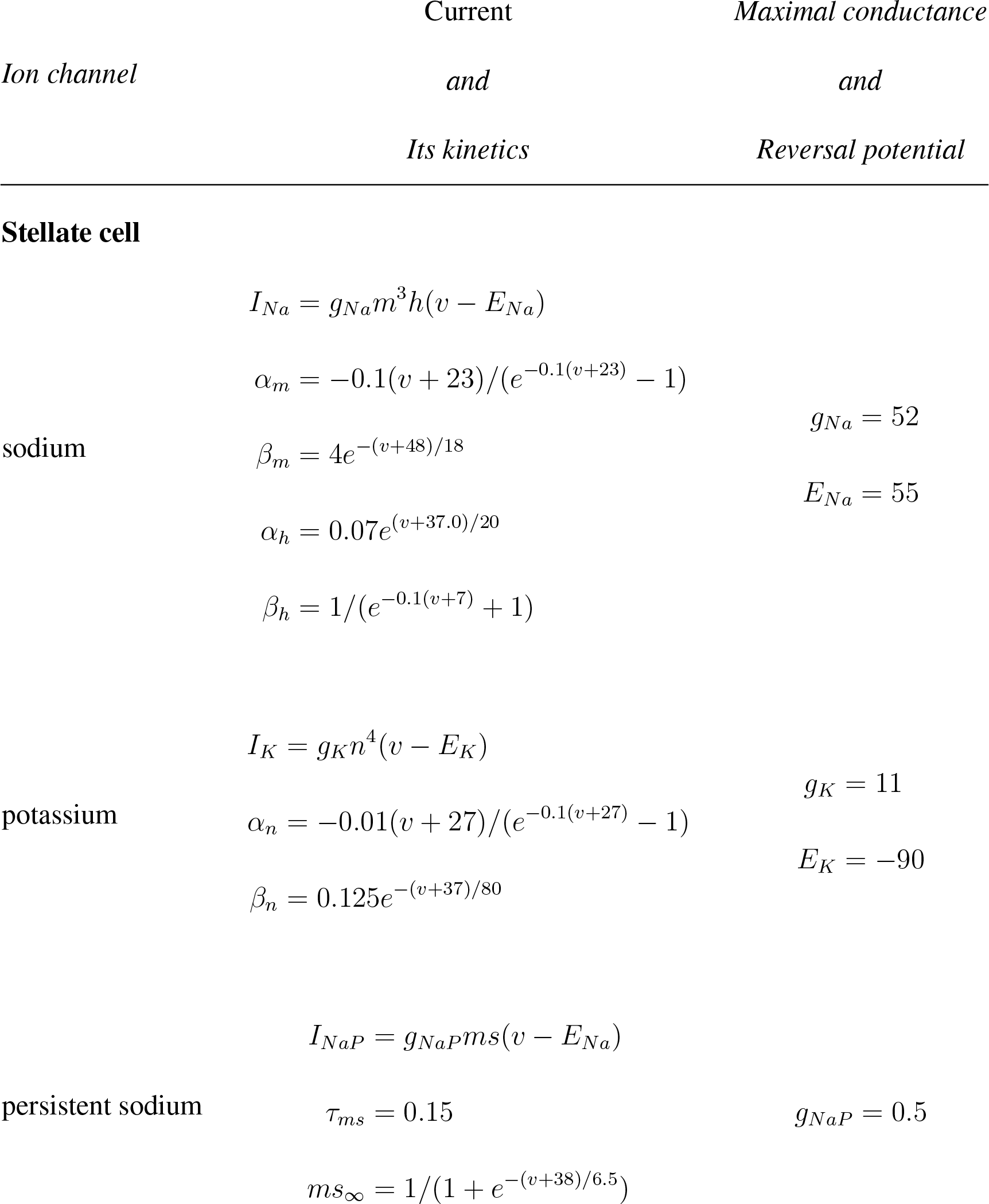
Functional form of conductances and gating variables

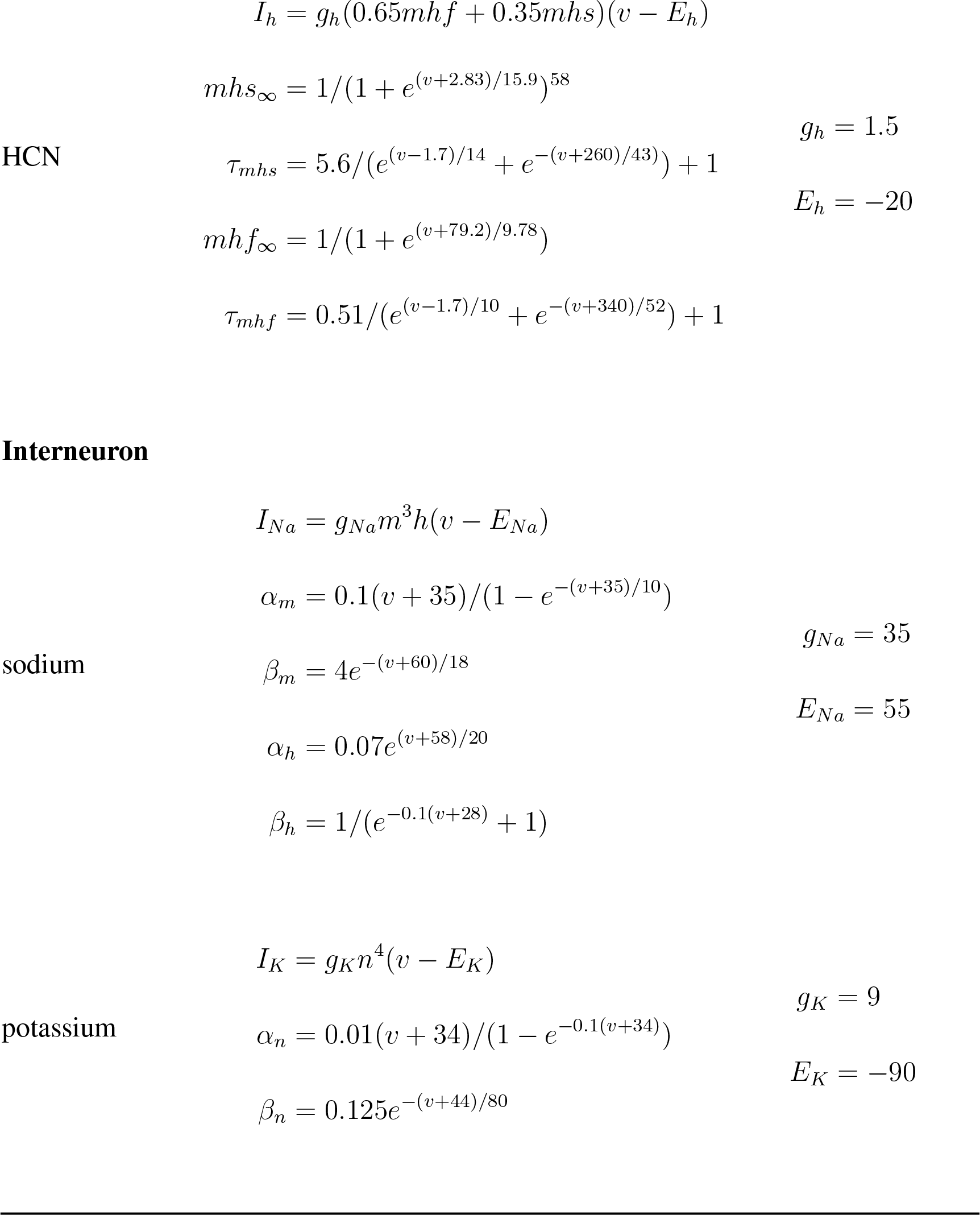

**Table 2:**
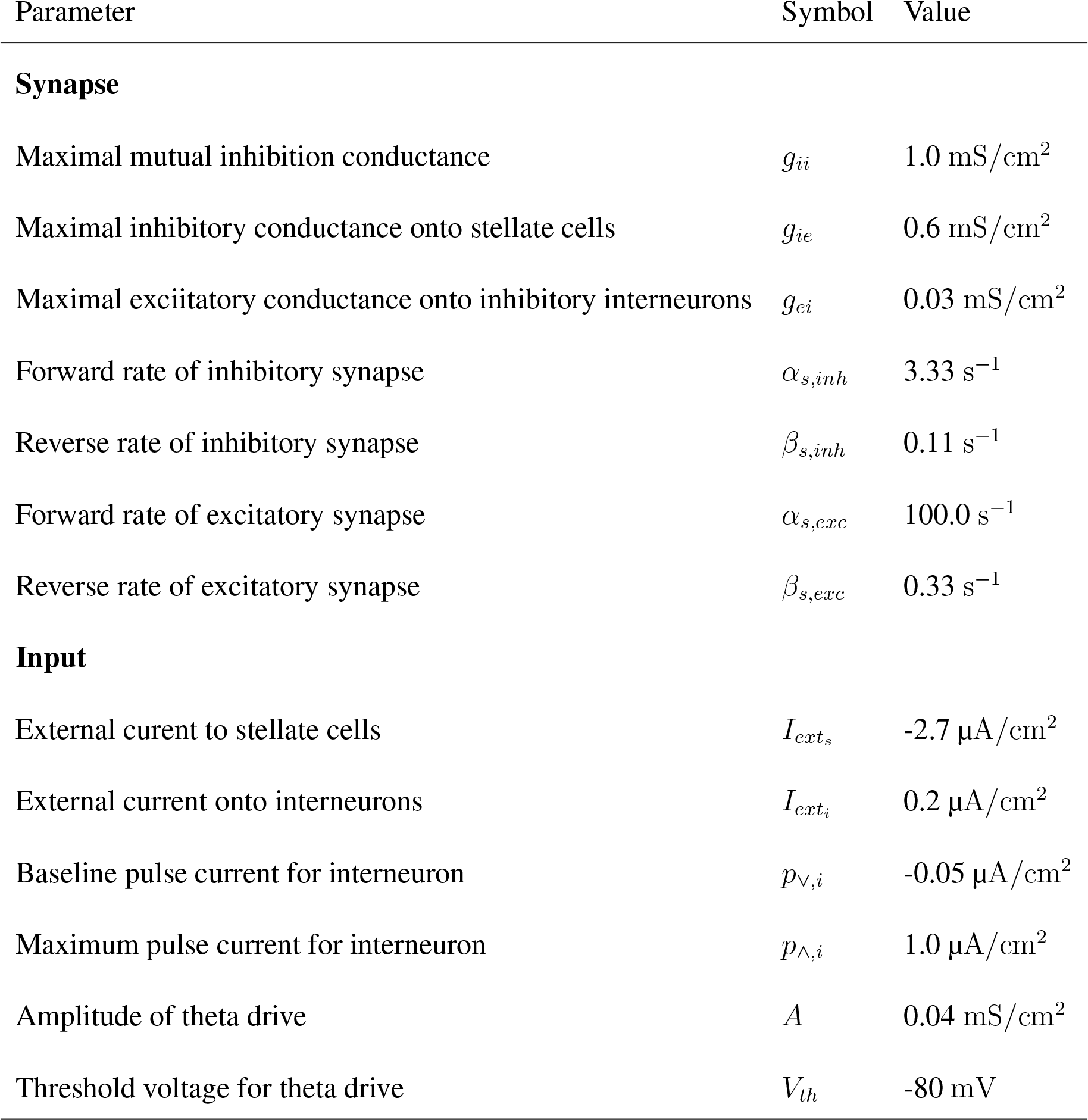
List of default network and input parameters

## Acknowledgements

CA was funded by DBT–Wellcome India Alliance through an Intermediate fellowship IA/I/11/2500290 and IISER Pune. AN was funded through a University Grants Commission Senior Research Fellowship and IISER Pune. We thank members of the Nadkarni and Assisi labs and Dr. Aurnab Ghose for useful discussions.

## References

1. Witter, M. P., Doan, T. P., Jacobsen, B., Nilssen, E. S. & Ohara, S. Architecture of the entorhinal cortex: a review of entorhinal anatomy in rodents with some comparative notes. Frontiers in Systems Neuroscience 11, 1–12 (2017).

2. Cappaert, N. L. M., Van Strien, N. M. & Witter, M. P. Hippocampal Formation. 511–573 (Academic Press, San Diego, 2015).

3. Hafting, T., Fyhn, M., Molden, S., Moser, M.-B. & Moser, E. I. Microstructure of a spatial map in the entorhinal cortex. Nature 436, 801–6 (2005).

4. Sargolini, F. et al. Conjunctive representation of position, direction, and velocity in entorhinal cortex. Science 758–763 (2006).

5. Solstad, T., Boccara, C. N., Kropff, E., Moser, M.-b. & Moser, E. I. Representation of geometric borders in the entorhinal cortex. Science 1109, 1865–1869 (2008).

6. Vertes, R. P. & Kocsis, B. Brainstem – diencephalo-septohippocampal systems controlling the theta rhythm of the hippocampus. Neuroscience 81, 893–926 (1997).

7. Koenig, J., Linder, A. N., Leutgeb, J. K. & Leutgeb, S. The spatial periodicity of grid cells. Science 592, 592–595 (2011).

8. Bolding, K. A., Ferbinteanu, J., Fox, S. E. & Muller, R. U. Place cell firing cannot support navigation without intact septal circuits. bioRxiv 470088 (2018).

9. Benchenane, K. et al. Coherent theta oscillations and reorganization of spike timing in the hippocampal-prefrontal network upon learning. Neuron 66, 921–936 (2010).

10. Takehara-Nishiuchi, K., Maal-Bared, G. & Morrissey, M. D. Increased entorhinal–prefrontal theta synchronization parallels decreased entorhinal–hippocampal theta synchronization during learning and consolidation of associative Memory. Frontiers in Behavioral Neuroscience 5, 1–13 (2012).

11. Wilson, D. I. G. et al. Lateral entorhinal cortex is critical for novel object-context recognition 366, 352–366 (2013).

12. Tsao, A. et al. Integrating time from experience in the lateral entorhinal cortex. Nature 561, 57–62 (2018). arXiv:0806.3143v1.

13. Gonzalez-Sulser, A. et al. GABAergic projections from the medial septum selectively inhibit interneurons in the medial entorhinal cortex. The Journal of Neuroscience 34, 16739 LP – 16743 (2014).

14. Nilssen, X. E. S. et al. Inhibitory connectivity dominates the fan cell network in layer II of lateral entorhinal cortex. Journal of Neuroscience 38, 9712–9727 (2018).

15. Couey, J. J. et al. Recurrent inhibitory circuitry as a mechanism for grid formation. Nature Neuroscience 16, 318–324 (2013).

16. Buetfering, C., Allen, K. & Monyer, H. Parvalbumin interneurons provide grid cell–driven recurrent inhibition in the medial entorhinal cortex. Nature Neuroscience 17, 710–718 (2014).

17. Fuchs, E. C. et al. Local and distant input controlling excitation in layer II of the medial entorhinal cortex. Neuron 89, 194–208 (2016).

18. Acker, C. D., Kopell, N. & White, J. a. Synchronization of strongly coupled excitatory neurons: Relating network behavior to biophysics. Journal of Computational Neuroscience 15, 71–90 (2003).

19. Rotstein, H. G., Oppermann, T., White, J. a. & Kopell, N. The dynamic structure underlying subthreshold oscillatory activity and the onset of spikes in a model of medial entorhinal cortex stellate cells. Journal of Computational Neuroscience 21, 271–292 (2006).

20. Wang, X.-j. Gamma oscillation by synaptic inhibition in a hippocampal interneuronal network model. Journal of Neuroscience 16, 6402–6413 (1996).

21. Alonso, A. & Klink, R. Differential electroresponsiveness of stellate and pyramidal-like cells of medial entorhinal cortex layer II. Journal of Neurophysiology 70, 128–143 (1993).

22. Dickson, C. T. et al. Properties and role of I(h) in the pacing of subthreshold oscillations in entorhinal cortex layer II neurons. Journal of Neurophysiology 83, 2562–2579 (2000).

23. Shay, C. F., Ferrante, M., Chapman 4th, G. W. & Hasselmo, M. E. Rebound spiking in layer II medial entorhinal cortex stellate cells: Possible mechanism of grid cell function. Neurobiology of learning and memory 129, 83–98 (2016).

24. Magistretti, J. & Alonso, A. Fine gating properties of channels responsible for persistent sodium current generation in entorhinal cortex neurons. The Journal of General Physiology 120, 855–873 (2002).

25. Ray, S. et al. Grid-layout and theta-modulation of layer 2 pyramidal neurons in medial entorhinal cortex. Science 1243028 (2014).

26. Rowland, D. C. et al. Functional properties of stellate cells in medial entorhinal cortex layer II. eLife 7, 1–17 (2018).

27. Krupic, J., Bauza, M., Burton, S., Barry, C. & O’Keefe, J. Grid cell symmetry is shaped by environmental geometry. Nature 518, 232–235 (2015).

28. Barry, C., Ginzberg, L. L., O’Keefe, J. & Burgess, N. Grid cell firing patterns signal environmental novelty by expansion. Proceedings of the National Academy of Sciences 109, 17687–17692 (2012). arXiv:1408.1149.

29. Wernle, T. et al. Integration of grid maps in merged environments. Nature Neuroscience 21, 92–105 (2018).

30. Shilnikov, A. L. & Maurer, A. P. The art of grid fields: geometry of neuronal time. Frontiers in Neural Circuits 10, 1–16 (2016).

31. Yoon, K. et al. Grid cell Responses in 1D environments assessed as slices through a 2D Lattice. Neuron 89, 1086–1099 (2016).

32. Jeewajee, A., Barry, C., O’Keefe, J. & Burgess, N. Grid cells and theta as oscillatory interference: Electrophysiological data from freely moving rats. Hippocampus 18, 1175–1185 (2008). arXiv:1011.1669v3.

33. Yartsev, M. M., Witter, M. P. & Ulanovsky, N. Grid cells without theta oscillations in the entorhinal cortex of bats. Nature 479, 103–107 (2011).

34. Akam, T. & Kullmann, D. M. Oscillations and filtering networks support flexible routing of information. Neuron 67, 308–320 (2010).

35. Akam, T. & Kullmann, D. M. Oscillatory multiplexing of population codes for selective communication in the mammalian brain. Nature Reviews Neuroscience 15, 111–122 (2014).

36. Eliav, T. et al. Nonoscillatory phase coding and synchronization in the bat hippocampal formation. Cell 175, 1119–1130 (2018).

37. Kay, L. M. Theta oscillations and sensorimotor performance. Proceedings of the National Academy of Sciences 102, 3863–3868 (2005). https://www.pnas.org/content/102/10/3863.ful.pdf.

38. Kim, J. & Lee, I. Neural Correlates of Object-in-Place Learning in Hippocampus and Prefrontal Cortex. Journal of Neuroscience 31, 16991–17006 (2011).

39. Liebe, S., Hoerzer, G. M., Logothetis, N. K. & Rainer, G. Theta coupling between V4 and prefrontal cortex predicts visual short-term memory performance. Nature Neuroscience 15, 456–462 (2012).

40. Fries, P. Rhythms for cognition: communication through coherence. Neuron 88, 220–235 (2016).

41. Adhikari, A., Topiwala, M. A. & Gordon, J. A. Synchronized activity between the ventral hippocampus and the medial prefrontal cortex during anxiety. Neuron 65, 257–269 (2010).

42. Jones, M. W. & Wilson, M. A. Theta rhythms coordinate hippocampal–prefrontal interactions in a spatial memory task. PLOS Biology 3 (2005).

43. Reinhart, R. M. G., Zhu, J., Park, S. & Woodman, G. F. Synchronizing theta oscillations with direct-current stimulation strengthens adaptive control in the human brain. Proceedings of the National Academy of Sciences 112 (2015).

44. Haas, J. S., Nowotny, T. & Abarbanel, H. Spike-timing-dependent plasticity of inhibitory synapses in the entorhinal cortex. Journal of Neurophysiology 96, 3305–3313 (2006). 08234.1712/

45. Mehta, M. R., Barnes, C. A. & McNaughton, B. L. Experience-dependent, asymmetric expansion of hippocampal place fields. Proceedings of the National Academy of Sciences 94, 8918–8921 (1997).

46. Ólafsdóttir, H. F., Bush, D. & Barry, C. The role of hippocampal replay in memory and planning. Current Biology 28, R37 – R50 (2018).

47. Kreuz, T., Chicharro, D., Houghton, C., Andrzejak, R. G. & Mormann, F. Monitoring spike train synchrony. Journal of Neurophysiology 109, 1457–1472 (2013).

48. Mulansky, M. & Kreuz, T. PySpike—A Python library for analyzing spike train synchrony.SoftwareX 5, 183–189 (2016).

